# TooManyCellsInteractive: a visualization tool for dynamic exploration of single-cell data

**DOI:** 10.1101/2023.06.16.544954

**Authors:** Conor Klamann, Christie Lau, Gregory W. Schwartz

## Abstract

As single-cell sequencing data sets grow in size, visualizations of large cellular populations become difficult to parse and require extensive processing to identify subpopulations of cells. Managing many of these charts is laborious for technical users and unintuitive for non-technical users. To address this issue, we developed TooManyCellsInteractive (TMCI), a browser-based JavaScript application for visualizing hierarchical cellular populations as an interactive radial tree. TMCI allows users to explore, filter, and manipulate hierarchical data structures through an intuitive interface while also enabling batch export of high-quality custom graphics. Here we describe the software architecture and illustrate how TMCI has identified unique survival pathways among drug-tolerant persister cells in a pan-cancer analysis. TMCI will help guide increasingly large data visualizations and facilitate multi-resolution data exploration in a user-friendly way.

## Introduction

Single-cell sequencing enables transcriptome and epigenome quantification at the resolution of individual cells. Such high-throughput technologies enable remarkable opportunities to characterize the cellular landscape of biological processes and disease. However, the increased complexity of single-cell data presents new challenges for effectively exploring and visually interpreting large numbers of cells.

While current tools for single-cell visualization have led to important biological findings, these bioinformatic workflows generate complex images when applied to large scale data sets. A key component of existing analytical workflows involves partitioning cells into distinct groups of similar sequencing profiles, or “cell states”^1–4^. This action often relies on clustering algorithms such as k-means, Louvain, or Leiden, all of which identify a single partitioning of cell states that is contingent on user-defined parameters and limited to a single clustering resolution^5^. To visualize cell states, workflows often incorporate dimensionality reduction to project cells, which regularly encompass over 10,000 features or dimensions such as genes or regions of DNA, to a two-dimensional surface using principal component analysis, t-distributed stochastic neighbor embedding (t-SNE), or uniform manifold approximation and projection (UMAP)^6–8^. However, the conventional approach of using scatter plot visualizations may not effectively communicate the structure of large data sets due to the high density of points on the plot. Scatter plots, due to their low dimensional representation, also cannot fully capture the true high-dimensional relationships between cells. As such, the distance between points on the scatter plot may not be informative^9, 10^ because not all cells arranged closer together exhibit similar cell states^11–13^. To overcome these limitations, we developed tools such as TooManyCells^1^ for single-cell RNA sequencing (scRNA-seq) data measuring gene expression and TooManyPeaks^2^ for single-cell sequencing assay for transposase-accessible chromatin (scATAC-seq) data measuring chromatin accessibility. These tools generate tree visualizations depicting the hierarchical relationships between large numbers of cells. Even with these tools, large trees must be manually pruned (collapsing children nodes into parent nodes) in order to increase interpretability. Such challenges of interpreting and communicating biological insights will only grow over time as sequencing data sets continue to increase in size.

In addition to the capacity constraints of existing workflows, many of these bioinformatic tools require computational expertise that is out of reach for non-technical users yet also demand biological insight for data exploration. To help bridge this gap, a few interactive tools exist to visualize and facilitate exploration of high-throughput single-cell data. Some of the most widely-accessible tools include CELLxGENE^14^ and Cirrocumulus^15^, among others^16, 17^. More recent tools have been designed to assist with specific challenges within the analytical workflow such as read alignment^18^, choosing clustering parameters^19^, intensive requirements for computational resources^20^, cell type annotation^21^, and lack of familiarity with shell scripting^22, 23^. However, their fundamental approaches towards single-cell data analysis remains unchanged from conventional workflows and do not address scalability issues.

To address these limitations, we introduce TooManyCellsInteractive (TMCI), a browser-based JavaScript application for visualizing cell clusters as an interactive radial tree. With its unique graphical user interface, TMCI can explore large single-cell data sets in an interactive way, revealing specific cellular populations in the tree. With TMCI, users can easily toggle display elements, drag nodes, re-color scale ranges, add gene expression overlays, prune nodes manually or by a variety of node properties, and export images and image specifications, all with an interactive interface. In addition to rendering the tree itself, TMCI provides a feature-rich dashboard for displaying metadata, including node counts and distributions, as well as pruning tools and a breadcrumb-style history bar. Here, we benchmark and compare TMCI with other commonly-used visualization tools across several data sets as well as demonstrate how TMCI can be used to delineate populations by applying TMCI on a data set of multiple cancer cell lines undergoing acute drug treatment. With the unique capabilities of TMCI, we report several commonly-used drug-resistance pathways across cancer types and identify a potential time-dependent hibernation state associated with drug tolerance. With an intuitive interface and flexible export system with batch processing options, TMCI is a one-stop solution to visualize large single-cell data sets. TMCI is open source and packaged with all dependencies at https://github.com/schwartzlab-methods/too-many-cells-interactive.

## Results

### Overview of TMCI usage and features

#### Inputs

TMCI consists of a browser-based graphical user interface (Figure 1), a web server, a relational database, a containerized runtime environment, and a collection of initialization and data processing scripts (Figure 2). TMCI requires two input files: a JSON representation of a tree and a CSV file that maps categorical labels to observation identifiers. Typically, these files are generated by the command-line TooManyCells application and will be named “cluster_tree.json” and “labels.csv”^1^. However, TMCI is compatible with input generated from alternative tools if each file has the appropriate structure. Optionally, the user may also provide feature data such as gene expression for the browser application to fetch and display as an overlay on the main visualization. We provide a convenience script “start-and-load.sh” as the main point of entry for the user, which will handle the database importation of Matrix Market Exchange (MEX) files or a directory of MEX files in compressed or uncompressed format through the “importMatrix.ts” helper script. For a step-by-step guide to feature data importation and viewing, we provide a visual tutorial in the project documentation located at https://schwartzlab-methods.github.io/too-many-cells-interactive/tutorial.html.

**Figure 1:**
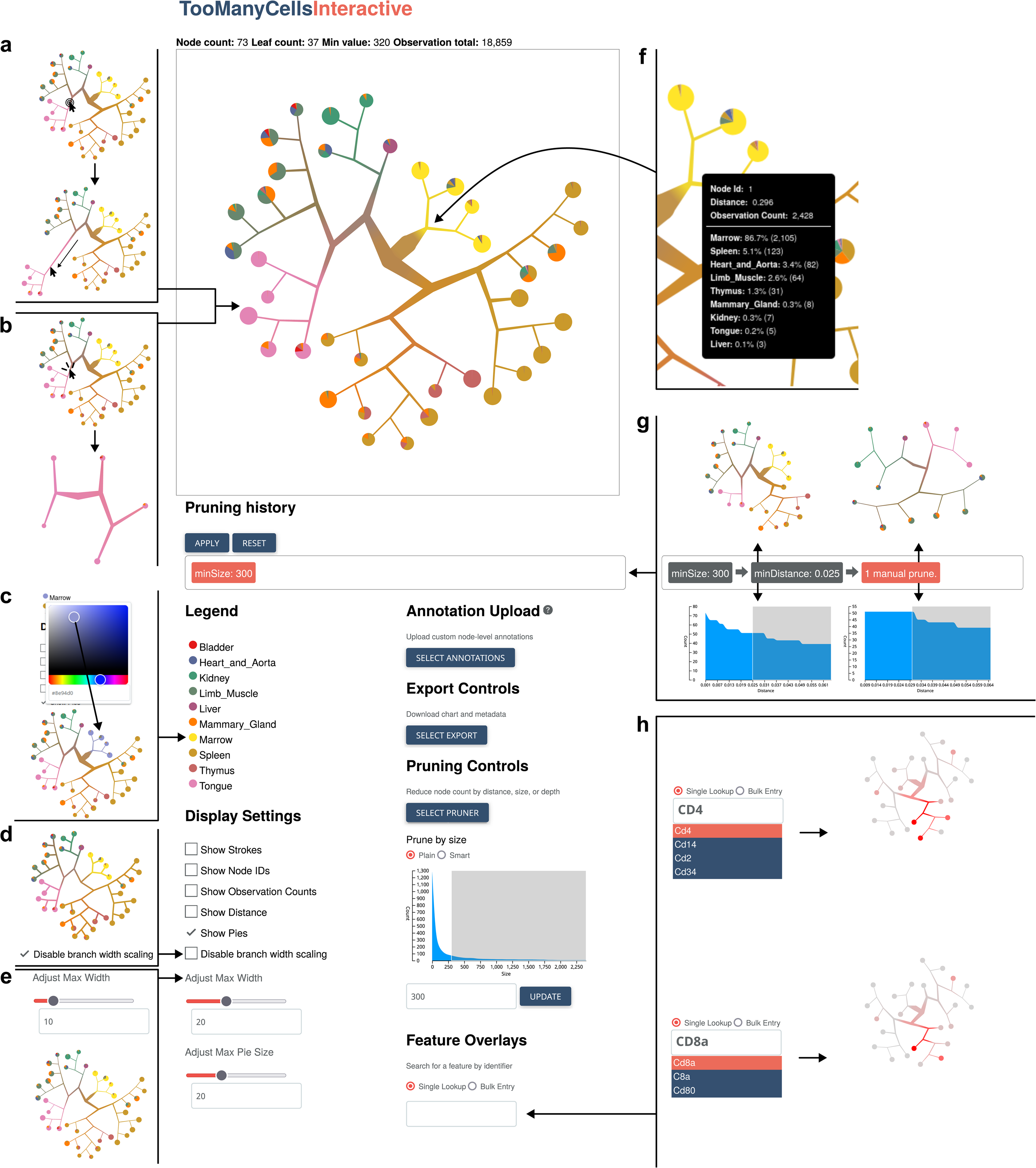
Overview of the TMCI output interface. (a,b) Direct interactions with the main interface. The user may manually edit the tree in the main interface by stretching or shrinking branches (a) or selecting a new tree root (b). (c) Color picker for cell labels through hex value or slider when selecting the label of choice in the legend. (d,e) Visualization features of the tree branches and nodes, including the disabling of branch scaling (d) and adjusting the width of branches (e), among other visualization features. (f) Live-updating tooltips containing statistics for each node. (g) Breadcrumb toolbar containing previous structural changes as the user interactively prunes the tree based on the distribution of nodes. (h) Fuzzy-search bar to see the overlay of a feature on each node in the tree, such as gene expression for each cellular population. The user may select one or several features through the fuzzy-search bar and select thresholds for “high” and “low” cutoffs for simultaneous feature overlays (e.g. both *CD4* and *CD8a*).

**Figure 2:**
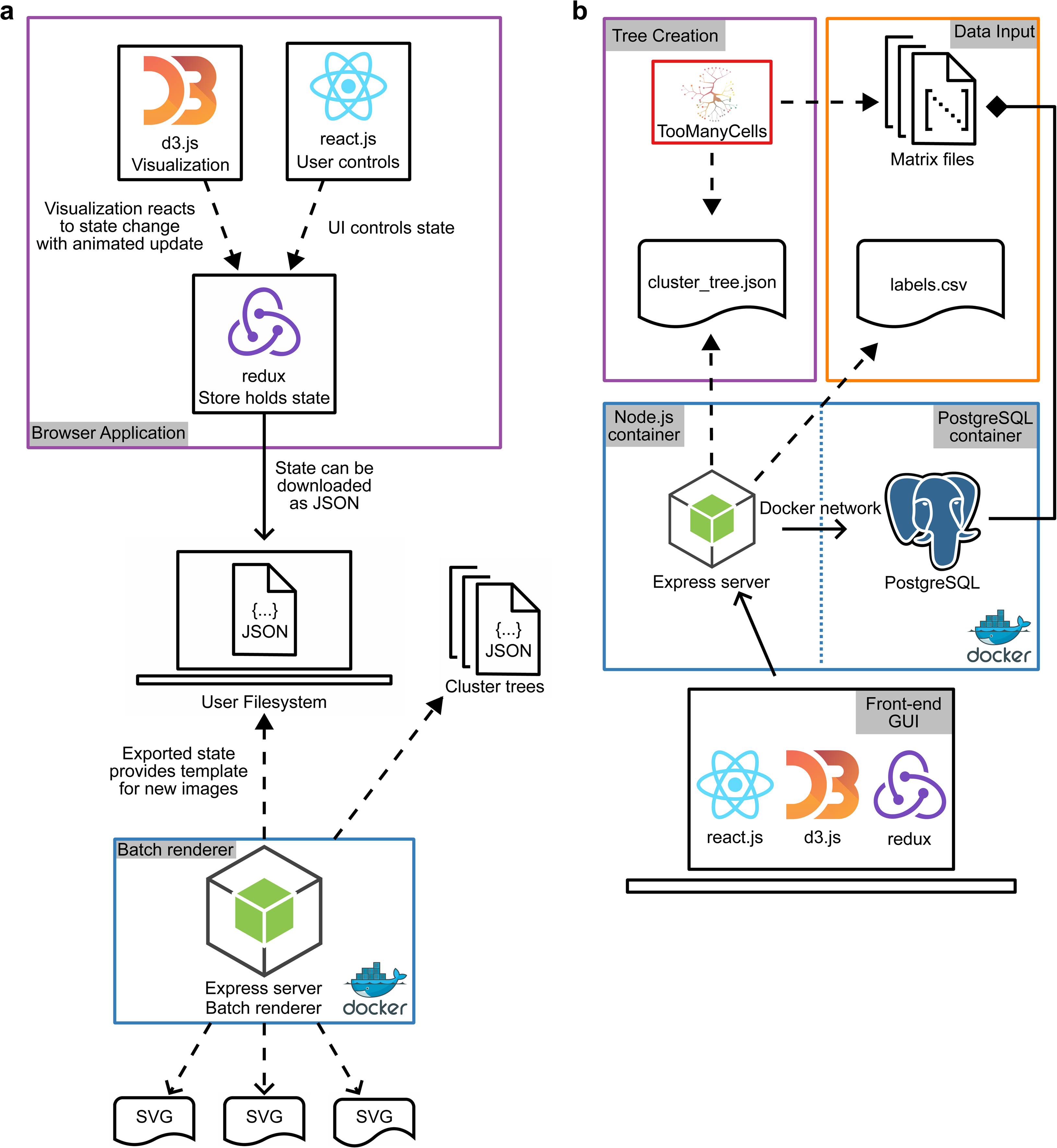
The architecture for TMCI. (a) The front-end architecture of TMCI. The user interacts with the D3 tree visualization and React user interface which sends state-change requests to Redux. This state-tracking feature enables batch processing: the user may upload a configuration state which the Express server will read without loading the graphical user interface and automatically export the corresponding SVG. (b) The back-end Express server container takes as input the tree structure and cell label files in the Node application. Similarly, a PostgreSQL container reads the matrix files containing the count matrices with features such as gene expression or chromatin accessibility. The Express container manages feature overlays on the tree through PostgreSQL queries in response to front-end requests. Flow charts are Unified Modeling Language structured diagrams where dashed closed arrows indicate dependencies, dashed open arrows indicate artifacts, solid closed arrows indicate relationships, and solid diamonds are compositions.

#### User Interface

TMCI offers both a headless, command-line interface for batch processing as well as a robust, easy-to-use graphical user interface with numerous features to best explore single-cell data.

#### Metadata summary

Situated above the main visualization, the metadata summary panel includes counts of visible nodes and leaves, the smallest value (node size) in the plot, and the number of cells for the entire data set.

#### Tree panel

The tree panel is an SVG DOM element that spans most of the left side of the viewport and frames the main visualization. Zoom, pan, and dragging behaviors are enabled in this area.

#### Radial tree visualization

Representing the user’s input as a radial tree, this interactive plot is the application’s principal visualization. Hovering over a node will display a tooltip with additional metadata and statistics, and nodes may be manipulated directly in several ways: they can be repositioned by dragging, collapsed by shift-clicking, or set as a new tree root by control-clicking.

#### Pruning history panel

This breadcrumb-style element at the top of the dashboard’s right-hand side or “control panel” represents the user’s tree pruning history. Each time users click the “Apply” button, their current pruning settings will be saved to the history panel as a discrete step. Clicking on a previous step will return the visualization to the corresponding previous state, while clicking the “Reset” button will remove all steps and revert the visualization to the original structure.

#### Scale selection panel

Located beneath the pruning history panel, the scale selection panel allows users to change the way tree values are displayed. The default scale, “Labels”, is an ordinal scale that displays feature labels as colors rendered as weighted blends. If one or more feature overlays has been selected, such as multiple gene expressions, the user may view features on (1) an ordinal scale of “high” or “low” combinatorial coloring based on selected feature value thresholds (e.g. *CD4* high and *CD8* high as red, *CD4* high and *CD8* low as blue) or (2) a sequential two-color scale indicating the average value of selected features (e.g. the average of *CD4* and *CD8* values). If the user has uploaded custom node annotations with pre-defined node values, they may view such annotations on a sequential two-color scale ranging from the lowest to highest value.

#### Legend

The legend can be found under the scale selection panel and displays by default the labels provided by the user and their corresponding colors as represented in the main graphic. If a feature or user annotation scale is selected, the legend will display high and low counts as well as the color gradient associated with the range of values. The legend allows users to modify scale colors by clicking on the corresponding color swatches.

#### Display element toggles

Beneath the legend, the user can use checkboxes to change the visibility of various graphical elements, including shape outlines (strokes), node identification labels, observation count annotations, distance indicators such as modularity from TooManyCells, leaf-node pie charts representing label proportion, and the presence of branch-width and pie-radius scaling.

#### Scale controls

The sliders under the display element toggles allow the user to adjust the maximum pixel size of pie charts and tree branch widths. The saturation slider can be used to improve the visibility of continuous scales.

#### Annotation uploader

As part of the control panel, the user can upload custom node annotations as a CSV file to overlay values directly on top of each node. This feature enables the user to overlay external results, such as gene set enrichment analysis normalized enrichment scores^24^, on the tree.

#### Export button

TMCI can export the current visualization state to several formats. For visual reproductions, PNG and SVG options are provided. JSON and CSV representations of the nodes are also available, along with the option to export a pruned tree in the rose-tree format given by TooManyCells. Finally, the user may choose to download the image configuration as a JSON file that can then be passed to the “headless” script for batch-generation with options chosen from the user interface.

#### Pruning controls

This element allows users to select one of four “pruners” in order to collapse children nodes and reduce the size of the tree. Users may prune by minimum node size, minimum node depth, or a minimum distance such as network modularity.

#### Pruner distribution chart

Each pruner has a corresponding brushable area chart representing the visible node count along the range of possible pruning values. The user may change the pruning value by dragging the slider to the desired value or by entering a number in the input box. After each prune is “applied”, the distributions are recalculated and the area chart updated in real time.

#### Feature selector and threshold indicator

The final element in the right-hand column of the control panel allows the user to overlay feature values on the visualization and adjust the threshold level for the high/low ordinal color scale. In order to use this element, users will need to upload a count matrix in advance. The input box is an autocomplete element that returns “fuzzy” matches for feature names. Once the user selects a feature, the browser application will display a new tree with feature values represented by differently-colored nodes, by default using a sequential gradient scale. The user may select multiple features individually or request bulk searches that retrieve many features at once.

### TMCI reduces time to display trees

To compare the computational time and memory of our TMCI approach to data visualization from both our original, static implementation as well as other commonly used single-cell data exploration tools CELLxGENE^14^ and Cirrocumulus^15^, we developed five benchmarks for common single-cell analyses: displaying a tree, pruning a tree, overlaying a feature, overlaying five features in succession, and loading and displaying all single-cell data. We ran these benchmarks using 54,220 cells from a scRNA-seq data set of 11 samples across five cancer cell lines (Figure 3a,b), as well as 18,859 cells (“subset”; Figure 3c,d) and 41,668 cells (Figure 3e,f) from the Tabula Muris data set containing 10 mouse organs^25^.

**Figure 3:**
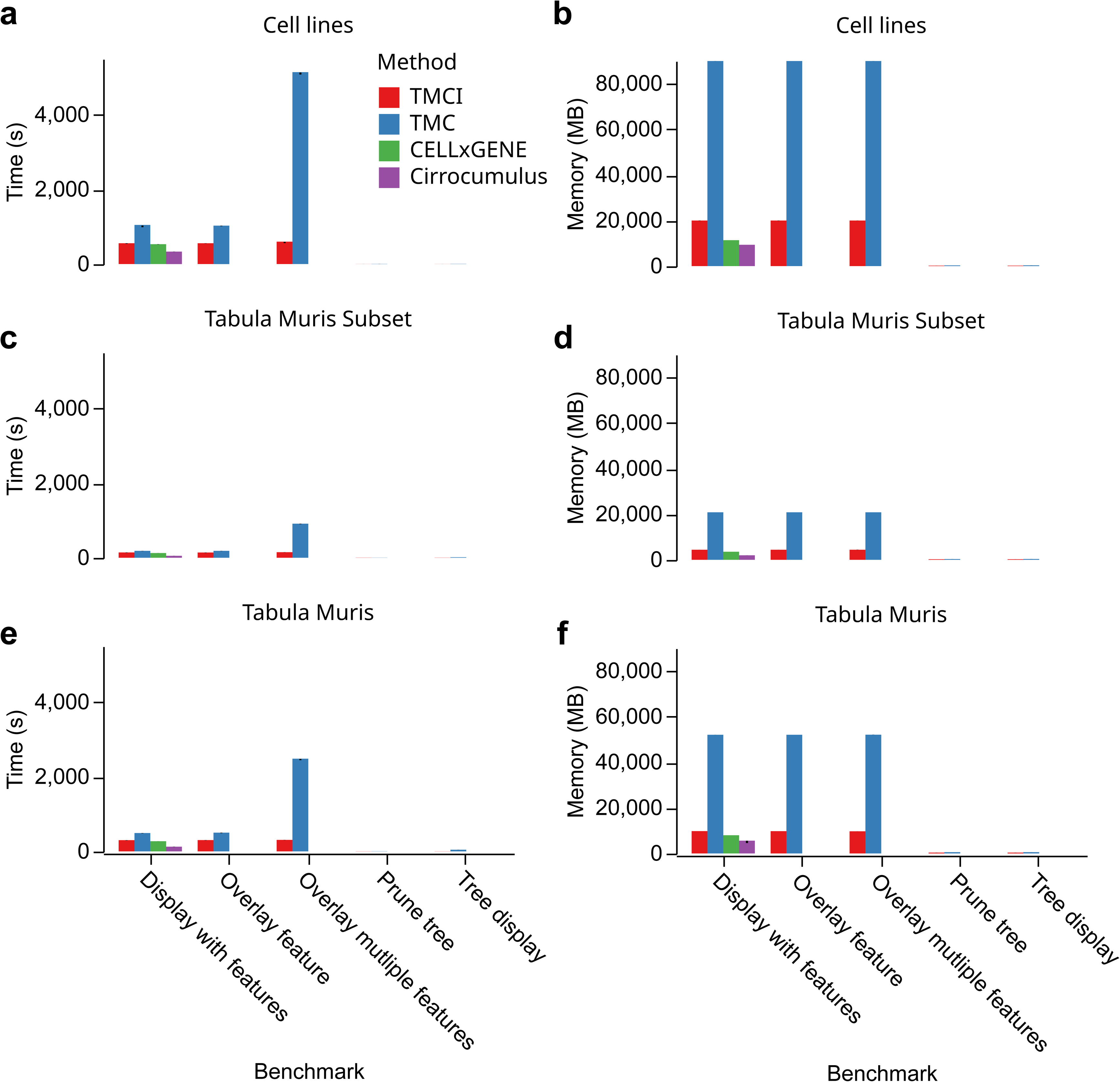
Comparative analysis of performance. (a-f) Comparisons on the *x*-axis including, from left to right, loading a count matrix and displaying a visualization (all programs), overlaying a single feature on a tree (tree programs only), batch processing five features on a tree (tree programs only), pruning a larger tree (tree programs only), and rendering a tree without the count matrix loaded (tree programs only). Comparisons were split by 11 samples from five cancer cell lines in response to drug treatment (a,b), a set of 10 samples from mouse tissues^25^ (c,d), or a set 24 samples from mouse tissues (e,f), measuring time (a,c,e) or memory usage (b,d,f). TMC: TooManyCells

To assess a baseline performance of each program, we compared the time and memory needed to display trees without feature overlays on our cancer cell line data set, meaning that no matrix processing was required. Inputs for both programs were the tree and label files generated by TooManyCells and we ran each benchmark five times to account for potential variability. TMCI was 4 fold faster than TooManyCells in the cancer cell line data set (mean 1.07 s vs. 4.62 s, *t*-test: *p* = 2.74 × 10^−18^; Figure 3a) demonstrating an order of magnitude speed improvement with our new implementation. Importantly, this upgrade did not come at the cost of memory, as TMCI used ∼ 120 MB less memory than TooManyCells (mean 188 MB vs. 308 MB, *t*-test: *p* = 1.87 × 10^−14^; Figure 3b). As this benchmark did not alter the structure of the tree, we next compared tools by pruning the tree to have nodes containing no fewer than 1,000 cells. This additional processing resulted in TMCI using approximately the same amount of resources as the unpruned tree and TooManyCells increasing its performance to a mean of 1.66 s (*t*-test: *p* = 2.39 × 10^−14^) and 300 MB of RAM (*t*-test: *p* = 5.56 × 10^−16^; Figure 3a,b). While the performance increase of TMCI over TooManyCells was consistent across data sets, some gains were 20 fold as with the larger Tabula Muris data set (Figure 3c-f, Supplementary Tables S1-S4).

Although TMCI outperformed with only tree display and processing, this benchmark did not account for matrix processing. As such, we next compared the performance of each program when rendering feature overlays, which introduces the resource-intensive task of retrieving expression data from the matrix. For a single feature on the cancer cell lines data set, TMCI outperformed TooManyCells in task duration (mean 554 s vs. 1,020 s, *t*-test: *p* = 1.91 × 10^−15^; Figure 3a). This advantage remained through TMCI’s greatly reduced memory usage (mean 19.9 GB vs. 89.8 GB, *t*-test: *p* = 2.11 × 10^−26^; Figure 3b). While this test displayed significant memory gains for TMCI over TooManyCells, a more applicable benchmark is to batch process the creation of several graphics from a single tree with varying gene expression overlays. In this benchmark of five features, TMCI outperformed TooManyCells in both time (mean 577 s vs. 5,100 s, *p* = 7.46 × 10^−18^) and memory (mean 19.9 GB vs. 89.8 GB, *p* = 2.44 × 10^−33^; Figure 3a,b) usage due to the unique persistent feature database, which enables TMCI to generate any number of images after only a single data import operation. TooManyCells, on the other hand, must process the matrix for each new graphic, leading to a linear 𝒪(*n*) performance where *n* is the number of feature overlays requested. As a result, TMCI was able to generate ten trees with just 23 s longer than a single tree, while TooManyCells took ten times longer than a single tree. These observations were consistent through all data sets (Figure 3c-f, Supplementary Tables S1-S4).

To compare with non-tree based methods, we measured the performance of loading an entire single-cell data set and producing a visualization. For our cancer cell line data set, Cirrocumulus was the fastest (mean 330 s), with CELLxGENE (mean 527 s) and TMCI (mean 550 s) close behind, and TooManyCells being the slowest (mean 1,010 s; Figure 3a). Likewise, Cirrocumulus had the lowest memory usage (mean 9.34 GB), followed by CELLxGENE (mean 11.3 GB), TMCI (mean 19.9 GB), and TooManyCells (mean 89.8 GB; Figure 3b). Importantly, TMCI displays all cluster resolutions, while CELLxGENE and Cirrocumulus only show “flat” clusterings, even though TMCI has closer performance to these two tools than TooManyCells which is significantly resource heavy (Supplementary Tables S1 and S2). These observations are consistent in the smaller Tabula Muris subset but not in the larger Tabula Muris data set, where TMCI has the second-lowest time and memory usage, outperforming CELLxGENE by three-fold time and four-fold memory (Figure 3e-f, and Supplementary Tables S1-S4). Together, these benchmarks indicate not only the superior performance of TMCI to a generate static and interactive tree of single-cell data compared to other tools, but also its ability to quickly and efficiently batch process many trees at once.

### Case study: TMCI effectively delineates subpopulations of cancer drug-tolerant persister cells

To demonstrate the utility of TMCI for quantification and visualization of relationships between diverse single-cell data sets, we explored the transcriptional differences induced by short-(2-3 days) and long-term (6-7 weeks) treatment of cancer cells *in vitro*. While treatment eliminates the majority of cancer cells, rare populations of drug-tolerant persister cells survive and may potentially act as a reservoir for drug-resistant growth^26^. Persister cells are characterized by a non-genetic, slow-cycling state that is reversible; upon drug holiday, persister cells are re-sensitized to treatment^27^. We sought to better understand the differences between short- and long-term treatment exposure in these persister cells using TMCI. To this end, we aggregated publicly available scRNA-seq data from five independent cancer persister-cell experiments across various disease areas and treatment modalities (Figure 4a and Table 1)^1, 28–31^. The TMCI visualization identified distinct separation between cancer cell lines, followed by division of control and treatment arms (Figure 4b). This hierarchy suggests that cells of a given cancer type, regardless of drug treatment, are more transcriptionally similar to one another than persister cells across cancer types for most populations.

**Figure 4:**
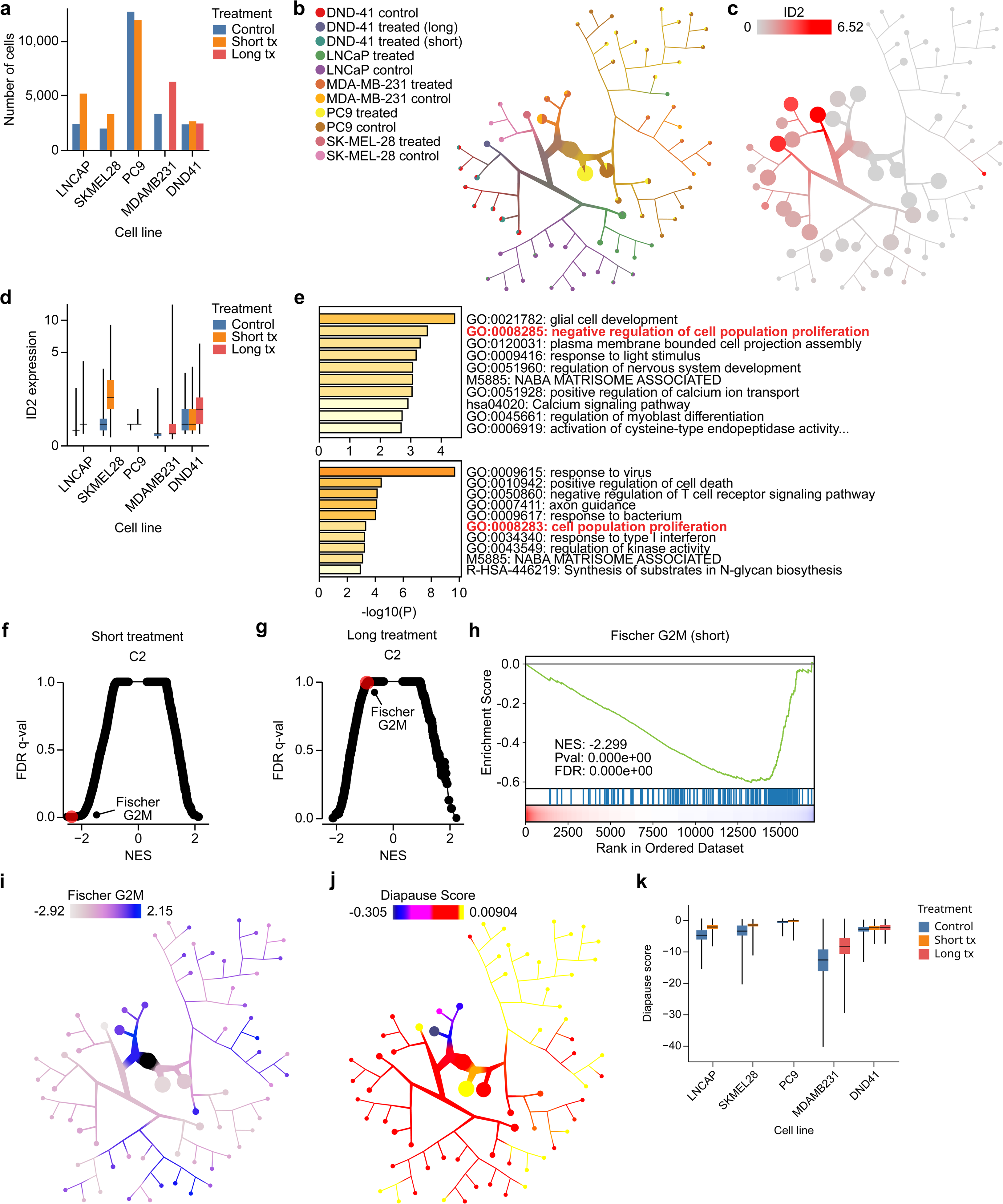
TMCI identifies distinct transcriptional programs across short-term and long-term treated drug-tolerant persister cells across cancer types. (a) Counts of pancreatic (LNCaP), melanoma (SK-MEL-28), lung (PC9), breast (MDA-MB-231) and T-cell acute lymphoblastic leukemia (DND-41) cancer cells from public persister-cell scRNA-seq experiments. Each cell line (untreated control: blue) received a short-(1-3 days: orange) or long-term (6-7 weeks: red) anti-cancer treatment. (b,c) TMCI tree of cells from (a) colored by cell line and treatment condition (b) or by expression of *ID2* (c). (d) Box-and-whisker plot of *ID2* expression for cells from each treatment condition (center line indicates median, box indicates interquartile range, whiskers indicate 1.5 ×interquartile range). (e) The top 10 enriched pathways determined by Metascape^38^ of the top 100 upregulated genes among short-term (top) and long-term (bottom) treated cells. Pathways relevant to the regulation of cellular proliferation are highlighted in red. (f,g) Normalized enrichment scores (NES) and respective *q*-values from gene set enrichment analysis (GSEA)^24^ of short-term (f) and long-term (g) treated cells compared to corresponding control cells. “FISCHER_G2_M_CELL_CYCLE” is highlighted in red. (h) GSEA curve of the “FISCHER_G2_M_CELL_CYCLE” gene set for short-term treated cells against untreated cells. (i) NES scores of the “FISCHER_G2_M_CELL_CYCLE” gene set for each node against all other nodes in the TMCI tree from (b). (j) TMCI tree from (b) colored by diapause gene signature. (k) Box-and-whisker of diapause signature scores for cells from each treatment condition.

**Table 1:**
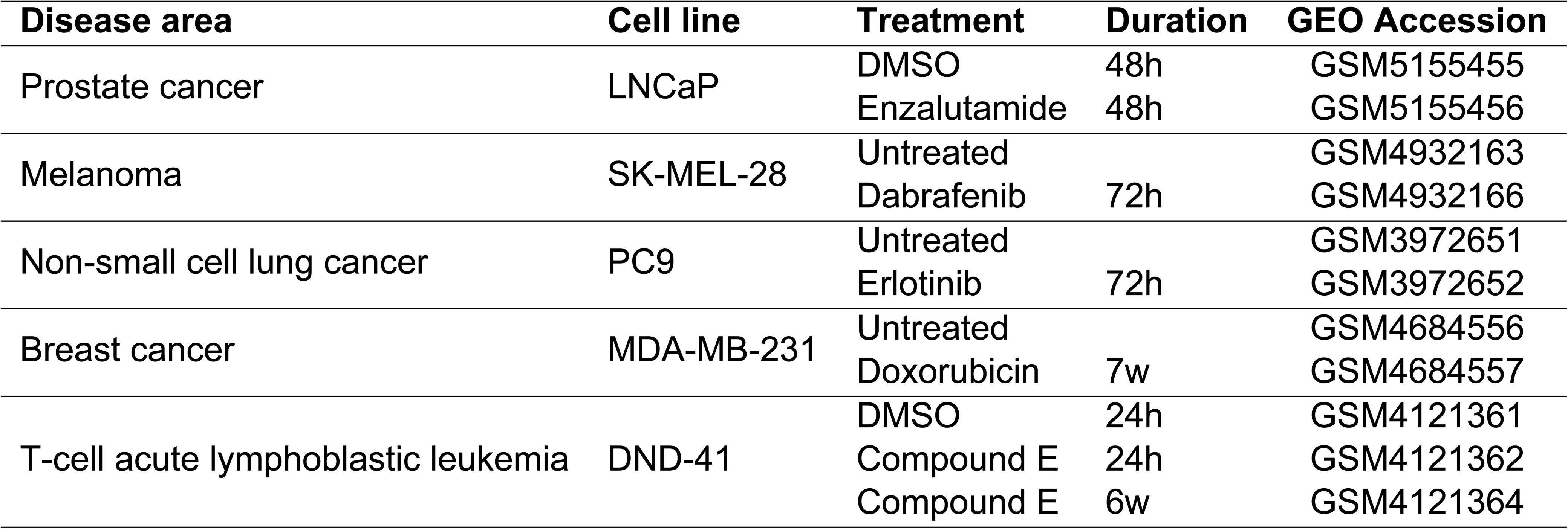
Human cancer cell lines from single-cell RNA-seqencing persister-cell experiments used in this case study. Corresponding anti-cancer drugs, treatment duration, and GEO accession numbers are listed.

### TMCI identified differentially expressed *ID2* across persister-cell populations

In order to understand how survival programs could be affected by the duration of treatment, we sought to characterize the unique expression profiles among persister-cell populations. The rank product of differential gene expression analysis of each cell line identified *ID2* as one of the most highly upregulated genes across long-term treated cells in comparison to controls (rank product: 4, permutation test: *p* < 2.22 × 10^−16^), but not among the short-term treated cells (rank product: 380, permutation test: *p* < 2.22 × 10^−16^). *ID2* is known to play a role in tumorigenesis as a key regulator of cell-cycle progression and overexpression of *ID2* in cell-line experiments modulates proliferative capacity and cell invasiveness^32, 33^. Visualization of *ID2* expression overlaid onto the tree showed enriched upregulation in cells within the long-term treated cell populations (Figure 4c). Comparison of *ID2* expression between each control and corresponding treatment arm showed a significant increase for all cell lines (Mann-Whitney *U* test: *p* < 0.05), regardless of treatment duration, with the exception of DND-41 which had higher *ID2* expression in long-term treatment (Figure 4d and Supplementary Tables S5-S7). This differential *ID2* expression suggests varying proliferative activity between treatment durations.

### TMCI identifies distinct proliferation mechanisms within persister-cell populations

To interrogate the ongoing biological mechanisms within short and long-term treated persister-cell populations, we performed pathway analysis using the top 100 upregulated differentially-expressed genes in the treated cells. Metascape^34^ analysis of the differentially-expressed genes from short-term treated cells identified “negative regulation of cell population proliferation” as a key biological process (hypergeometric test: *p* = 2.88 × 10^−4^; Figure 4e). Conversely, the same analysis performed on differentially-expressed genes identified from long-term treated cells suggest an increase of cellular proliferation across pathways (hypergeometric test: *p* = 4.89 × 10^−4^; Figure 4e). Subsequent exploration of the full list of differentially expressed genes using Gene Set Enrichment Analysis^24^ returned markedly distinct biological programs between the short and long-term treated populations. Among short-term treated populations, the most significantly decreased hits were found to be associated with various proliferation and cell-cycle-regulation programs. In line with our previous findings, these programs were not significantly downregulated among long-term treated cells (Supplementary Table S8). Among these gene sets, we found the expression of “FISCHER_G2_M_CELL_CYCLE” significantly decreased among short-term treated cells (NES = −2.30, Kolmogorov-Smirnov test: *p* < 2.22 × 10^−16^) but not among long-term treated cells (NES = −0.891, Kolmogorov-Smirnov test: *p* = 0.747; Figure 4f-i and Figure S1a). Consistent with this observation, additional G2M checkpoint and E2F target gene sets showed similar patterns (Supplementary Table S9). These findings suggest that persister cells utilize distinct pathways associated with modulation of proliferation and cell cycling throughout the duration of treatment.

### TMCI identifies subpopulations with highly expressed diapause programs

As we identified proliferation and cell-cycle factors associated with treatment duration, we were interested in understanding the temporal expression of diapause programs within the short- and long-term treatment persister-cell populations. Diapause is a reversible state of suspended embryonic development triggered by adverse environmental conditions^35^. Similarly, persister cells which survive throughout exposure to treatment undergo transcriptional adaptations resembling a diapause-like state^36, 37^. Overlaying diapause gene signature scores on the tree structure showed enrichment in all treated subpopulations compared to controls (Figure 4j and Figure S1b). Comparison between each control and treatment arm showed significantly increased diapause signature scores in all treated cell lines, again regardless of treatment duration (Mann-Whitney *U* test: *p* < 0.05; Figure 4k, Figure S1c-f, and Supplementary Table S10). For DND-41, which includes measurements of both short- and long-term treatment durations, the median diapause score increased from control to short-term to long-term, suggesting a direct correlation between diapause gene signature scores and treatment duration. Together, our analysis points to persister cells with different proliferation activity depending on treatment duration.

## Discussion

As high-throughput single-cell technologies continue to measure increasing numbers of cells, we need new visualization tools to better identify and interpret cell states. Here we present TMCI as a powerful, interactive solution that simplifies data exploration of large data sets. These visualizations are intuitive, supporting easy tree manipulation through statistical or manual pruning, color mapping, feature overlays, and more. With these features, identification of rare cellular populations is straightforward compared to previous iterations of single-cell data figures. Importantly, these benefits are not at the cost of performance, with TMCI either outperforming or on-par with alternative interactive visualizations. As we implemented TMCI as a web server, users can easily and quickly access large data sets with little computational impact on their local host. As TMCI is fast, its additional capabilities to batch process many trees enables quick plotting of thousands of trees, all informed by a configuration file exported from manual choices derived from a single tree.

Using the numerous features afforded by TMCI, we delineated cellular populations from drug-treated cancer cell lines and identified distinct transcriptional programs between short- and long-term treated cell lines. These programs included cell-proliferation pathways downregulated in short-term persister cell states which are then subsequently lost in the long-term cellular populations across all cancer types measured. This finding extended to the diapause signature, which was increased in persister cells, in concordance with previous studies, but here across cancer type. Together, TMCI identified transcriptional programs that are dependent on treatment duration, suggesting further investigation on the timing of treatment for persister cells. Through this demonstration, we show that big data visualization tools will be necessary as available data grows, and we provide TMCI as a one-stop solution for identifying tree-based relationships in such data.

## Data Availability

The GEO accession numbers for each data set reported in this paper are GSM5155455 and GSM5155456 (prostate cancer); GSM4932163 and GSM4932166 (melanoma); GSM3972651 and GSM3972652 (non-small cell lung cancer); GSM4684556 and GSM4684557 (breast cancer); GSM4121361, GSM4121362, and GSM4121364 (T-cell acute lymphoblastic leukemia).

## Code Availability

TMCI is available at https://github.com/schwartzlab-methods/too-many-cells-interactive, with a tutorial at https://schwartzlab-methods.github.io/too-many-cells-interactive/. TMCI will read any tree in the appropriate format, or the original TooManyCells command-line tool may create a tree, located at https://github.com/GregorySchwartz/too-many-cells. Code for analyses within this paper are available at https://github.com/schwartzlab-methods/ too-many-cells-interactive-paper-analyses.

## Authors Contributions

G. W. S. conceived and supervised the project. C. K. developed the tool and benchmarks. C. K. ran and analyzed benchmarks. C. L. collected, ran, and analyzed cancer cell line data. C. K., C. L., and G. W. S. wrote the manuscript.

## Methods

### Implementation

The TMCI browser application is written in TypeScript, a statically-typed superset of JavaScript, and implements a variety of frameworks and libraries to provide a highly interactive graphical user interface. Principal UI elements include an interactive radial tree for data visualization and a dashboard-style panel of input controls enabling users to make real-time adjustments to their plots. Such adjustments include node filtering (“pruning”), scale modification, feature overlay, and manual position adjustment. For saving the tree, TMCI supports image exporting to both PNG and SVG formats.

The browser application’s base architecture is provided by custom React.js components, while state management is handled by Redux and the interactive plots are created with D3.js, a widely-used low-level collection of data-visualization modules for scaling, event binding, DOM traversal, and high-performance animations (Figure 2).

The back-end Node application transpiles the Typescript to JavaScript using a Webpack bundler and serves it to the user’s browser via an Express application (Figure 2). If users wish to include custom feature overlays in their plots, such as gene expression data, they may upload the data to the PostgreSQL database that has been configured to connect to the Node server.

Both the PostgreSQL database and the Node server run in Docker containers, for which TMCI provides a declarative configuration via Docker Compose. TMCI’s containerized architecture allows it to be run on any computer with Docker installed, and the TMCI codebase includes Bash scripts intended as convenience wrappers around commonly-used Docker commands that can be easily extended for custom use.

Because D3.js has no strict browser dependencies, TMCI’s radial tree plots can be rendered without a browser interface. TMCI provides both a Node script and a shell script to enable easy programmatic rendering. The scripts require the necessary input files (“cluster_tree.json”, “labels.csv”) as well as a configuration JSON string that can be exported directly from the browser interface. Thus, users may refine their visualizations in the graphical environment and then re-use their configurations as templates for scripted batch processing on the server.

### Benchmarks

We performed benchmarks using an AWS EC2 instance running Ubuntu 20.04 and Docker 20.10.17 with 64x Intel Xeon Platinum 8375C CPU @ 2.90 GHz and 534 GB RAM. Each benchmark used two sets of the Tabula Muris^25^ data set, one of which contained 18,859 cells from 10 tissue samples and one of which contained 41,668 cells from 24 tissue samples, both across bladder, heart and aorta, kidney, limb muscle, liver, mammary gland, bone marrow, spleen, thymus, and tongue. We ran each benchmark five times to account for variability in processing time and memory.

### Preprocessing of drug-treated cancer scRNA-seq data

To demonstrate the utility of TMCI, we investigated drug-tolerant persister-cell populations which are capable of surviving anti-cancer drug treatment through non-genetic programming of reversible mechanisms^27^. To study these persister cells, we analyzed drug response scRNA-seq data across five cell lines of prostate cancer, line melanoma, non-small cell lung cancer, breast cancer, and T-cell acute lymphoblastic leukemia (Table 1). The duration of anti-cancer drug treatment for each cell lines varied from short-term (2–3 days) to long-term (6–7 weeks), enabling the identification of persister cells across cancer types and time.

We aggregated the raw count matrices into a single AnnData object^39^. For each cell, we calculated diapause expression scores from an experimentally-curated gene set^36^. We then used the aggregated data set as input for analysis in TooManyCells, which filtered cells based on the default parameters of a minimum of 250 transcripts per cell and genes based on having at least one cell expressing the gene. To normalize cells between data sets, we used term frequency-inverse document frequency to weigh genes such that more frequent genes across cells had less impact on downstream clustering analyses.

### Generating drug-treated cancer cell trees

Using our filtered and normalized count matrices, we used TooManyCells to generate a tree structure and identify transcriptionally distinct groups within our data set^1^. In brief, TooManyCells implements a matrix-free hierarchical spectral clustering approach^40^ to recursively partition scRNA-seq cell data into similar groups, represented as a tree structure, and uses Newman-Girvan modularity^41^ as an indicator for reaching a leaf in the tree. We used the resulting tree structure from TooManyCells as input for TMCI and conducted minimum distance search pruning at a cutoff of of 0.019 to improve the visibility of small sub-populations.

### Measuring differential expression across cellular populations

Using the pruned hierarchical tree structure, we performed differential gene expression analysis between persister and control cells from their predominant nodes for each cell line. Using the resulting log_2_ fold change gene lists for each comparison, we combined all differential gene expression lists into a single list using rank product to identify the top most differentially expressed genes from all as well as short- and long-term treatment experiments independently. To identify pathways associated with each comparison, we conducted gene set enrichment analysis^24^ to compare cells within each node of the tree structure against all other cells using the MSigDB Hallmark, C2 (curated) and C6 (oncogenic) gene sets^42^.

