## Supplementary Information for "TooManyCellsInteractive: a visualization tool for dynamic exploration of single-cell data"

### Supplementary Figures

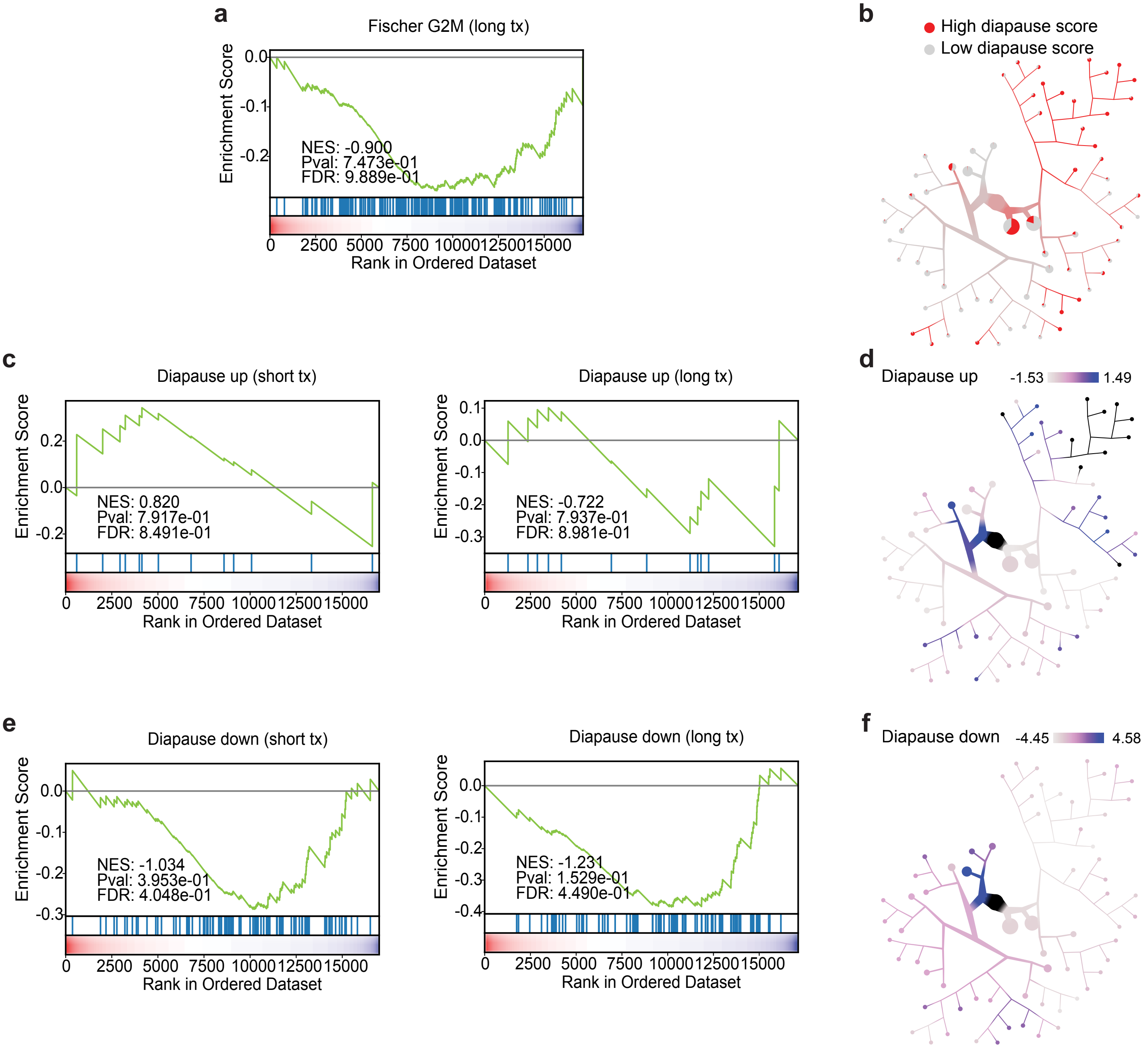
a
Fischer G2M (long tx)
Enrichment Score
NES: -0.900
Pval: 7.473e-01
FDR: 9.889e-01
Rank in Ordered Dataset
b
High diapause score
Low diapause score
c
Diapause up (short tx)
NES: 0.820
Pval: 7.917e-01
FDR: 8.491e-01
Diapause up (long tx)
NES: -0.722
Pval: 7.937e-01
FDR: 8.981e-01
d
Diapause up
-1.53
1.49
e
Diapause down (short tx)
NES: -1.034
Pval: 3.953e-01
FDR: 4.048e-01
Diapause down (long tx)
NES: -1.231
Pval: 1.529e-01
FDR: 4.490e-01
f
Diapause down
-4.45
4.58

Supplementary Figure S1: Short-term treatment across cell lines have lower proliferative enrichment. (a) Gene set enrichment analysis (GSEA) curve for the "FISCHER_G2_M_CELL_CYCLE" gene set in long-term treated cells. (b) TMCI tree from Figure 4b colored by diapause score (red: greater than 1 median absolute deviation away from the median diapause score; gray: otherwise). (c) GSEA curves of the upregulated diapause score gene set from short-term (left) or long-term (right) treated cells. (d) TMCI tree from Figure 4b colored by normalized enrichment scores the upregulated diapause gene set comparing each node to every other node. (e) GSEA curves of the downregulated diapause score gene set as in (c). (f) Normalized enrichment scores for the downregulated diapause score gene set as in (d).

### Supplementary Tables

#### Supplementary Table S1

Mean execution time for each performance benchmark across cells from 10 mouse tissues[25] ($n = 18{,}859$ cells), 24 mouse tissues ($n = 41{,}688$ cells), or 11 samples from five cancer cell lines in response to drug treatment ($n = 54{,}220$ cells).

#### Supplementary Table S2

Pairwise $t$-test results for each execution time performance benchmark across data sets from Supplementary Table S1. Benjamini–Hochberg method was used for testing and adjustment of $p$-values.

#### Supplementary Table S3

Mean memory usage for each performance benchmark across data sets from Supplementary Table S1.

#### Supplementary Table S4

Pairwise $t$-test results for each memory usage performance benchmark across data sets from Supplementary Table S1. Benjamini–Hochberg method was used for testing and adjustment of $p$-values.

#### Supplementary Table S5

Rank product of differentially expressed genes between long-term treated cells ($n = 8{,}654$) and their corresponding controls ($n = 5{,}639$). Mann-Whitney $U$ test was used to calculate $p$-values.

#### Supplementary Table S6

Rank product of differentially expressed genes between short-term treated cells ($n = 22{,}965$) and their corresponding controls ($n = 16{,}962$). Mann-Whitney $U$ test was used to calculate $p$-values.

### Supplementary Table S7

$\log_2$ fold change of gene expression between treated cells and their corresponding controls within a given cell line. Mann-Whitney *U* test was used to calculate *p*-values, Benjamini–Hochberg method was used for testing and adjustment of *p*-values.

### Supplementary Table S8

Gene set enrichment analysis results for long-term treated cells ($n = 8{,}654$) in comparison to their corresponding controls ($n = 5{,}639$), using the Hallmark, C2, and C6 gene sets.

### Supplementary Table S9

Gene set enrichment analysis results for short-term treated cells ($n = 22{,}965$) in comparison to their corresponding controls ($n = 16{,}962$), using the Hallmark, C2, and C6 gene sets.

### Supplementary Table S10

Comparison of diapause scores between treated cells and their corresponding control conditions within a given cell line. Mann-Whitney *U* test was used to calculate *p*-values, Benjamini–Hochberg method was used for testing and adjustment of *p*-values.
